# Cell Ecosystem and Signaling Pathways of Primary and Metastatic Pediatric Posterior Fossa Ependymoma

**DOI:** 10.1101/2020.08.10.244483

**Authors:** Rachael Aubin, Emma C. Troisi, Adam N. Alghalith, MacLean P. Nasrallah, Mariarita Santi, Pablo G. Camara

**Affiliations:** Department of Genetics and Institute for Biomedical Informatics, Perelman School of Medicine, University of Pennsylvania, Philadelphia, PA 19104, USA; Department of Pathology and Laboratory Medicine, Perelman School of Medicine, University of Pennsylvania, Philadelphia, PA 19104, USA; Department of Pathology, Children’s Hospital of Philadelphia, Philadelphia, PA 19104, USA

## Abstract

Pediatric ependymoma is a devastating brain cancer marked by its relapsing pattern and lack of effective chemotherapies. This shortage of treatments is partially due to limited knowledge about ependymoma tumorigenic mechanisms. Although there is evidence that ependymoma originates in radial glia, the specific pathways underlying the progression and metastasis of these tumors are unknown. By means of single-cell transcriptomics, immunofluorescence, and *in situ* hybridization, we show that the expression profile of tumor cells from pediatric ependymomas in the posterior fossa is consistent with an origin in LGR5+ stem cells. Tumor stem cells recapitulate the developmental lineages of radial glia in neurogenic niches, promote an inflammatory microenvironment in cooperation with microglia, and upon metastatic progression initiate a mesenchymal program driven by reactive gliosis and hypoxia-related genes. Our results uncover the cell ecosystem of pediatric posterior fossa ependymoma and identify WNT/β-catenin and TGF-β signaling as major drivers of tumorigenesis for this cancer.

## Introduction

Pediatric ependymoma remains a significant therapeutic challenge despite the general improvement of adjuvant chemotherapies for pediatric brain tumors during the past decades (Bouffet and Foreman, 1999; Bouffet et al., 1998; Duffner et al., 1998; Merchant et al., 2009). A roadblock to expanding treatment options is the current limited knowledge about the molecular mechanisms that underlie ependymoma tumorigenesis, progression, and metastasis. Genome-wide DNA-methylation profiling of ependymal tumors has led to their classification into nine groups associated with distinct anatomical location, age of onset, and prognosis (Mack et al., 2014; Pajtler et al., 2015; Witt et al., 2011). In young children, the most common group is posterior fossa ependymoma group A (PFA), representing approximately 70% of cases (Pajtler et al., 2015). In contrast with supratentorial ependymomas, PFA tumors have a balanced genome and do not have highly recurrent genetic alterations. Based on their methylation profile, PFA tumors can be further divided into two subgroups, PFA-1 and PFA-2, with similar clinical characteristics but distinct histogenesis (Pajtler et al., 2018). These subgroups express genes that are respectively active in the brainstem and the isthmic organizer during embryogenesis (Pajtler et al., 2018; Vladoiu et al., 2019), indicating a possible origin in radial glia from these locations (Johnson et al., 2010; Taylor et al., 2005). However, little is known about the molecular characteristics of ependymoma stem cells and the changes that lead to their metastasis into other regions of the central nervous system. To address this limitation, we have investigated the single-cell expression programs associated with the cell ecosystem, tumor-derived cell lineages, and stem cells of pediatric PFA ependymoma. By comparing the cell populations and gene expression programs between primary tumors and distal metastases, and relating them to neurodevelopmental and brain injury processes, we have identified some of the oncogenic pathways associated with the progression and metastases of these tumors.

## Results

### A transcriptomic atlas of primary and metastatic PFA ependymoma

To characterize the cell ecosystem of PFA tumors, we considered a cohort of 42 pediatric ependymal tumors collected and profiled for gene expression at the bulk level by the Children’s Brain Tumor Tissue Consortium (CBTTC). This cohort consisted of 37 primary tumors in the posterior fossa and 5 spinal or cortical metastases derived from posterior fossa primary tumors (Supplementary Table 1). We trained a classifier to disaggregate the cohort into known molecular groups based on the tumor gene expression profile (Methods). Consistent with other studies (Pajtler et al., 2018; Pajtler et al., 2015), most tumors in this cohort (40 out of 42) were classified as PFA, out of which 68% (*n* = 27) were identified as PFA-1 (Supplementary Table 1). We selected 6 primary and 3 metastatic PFA tumors from this cohort and performed massively parallel single-nuclei RNA sequencing (Hu et al., 2017) using archived flash-frozen tissue. Overall, we profiled the transcriptome of 26,724 cells and used these data to create a transcriptomic atlas of primary and metastatic PFA ependymoma (Figs. 1a and 3a).

**Figure 1.**
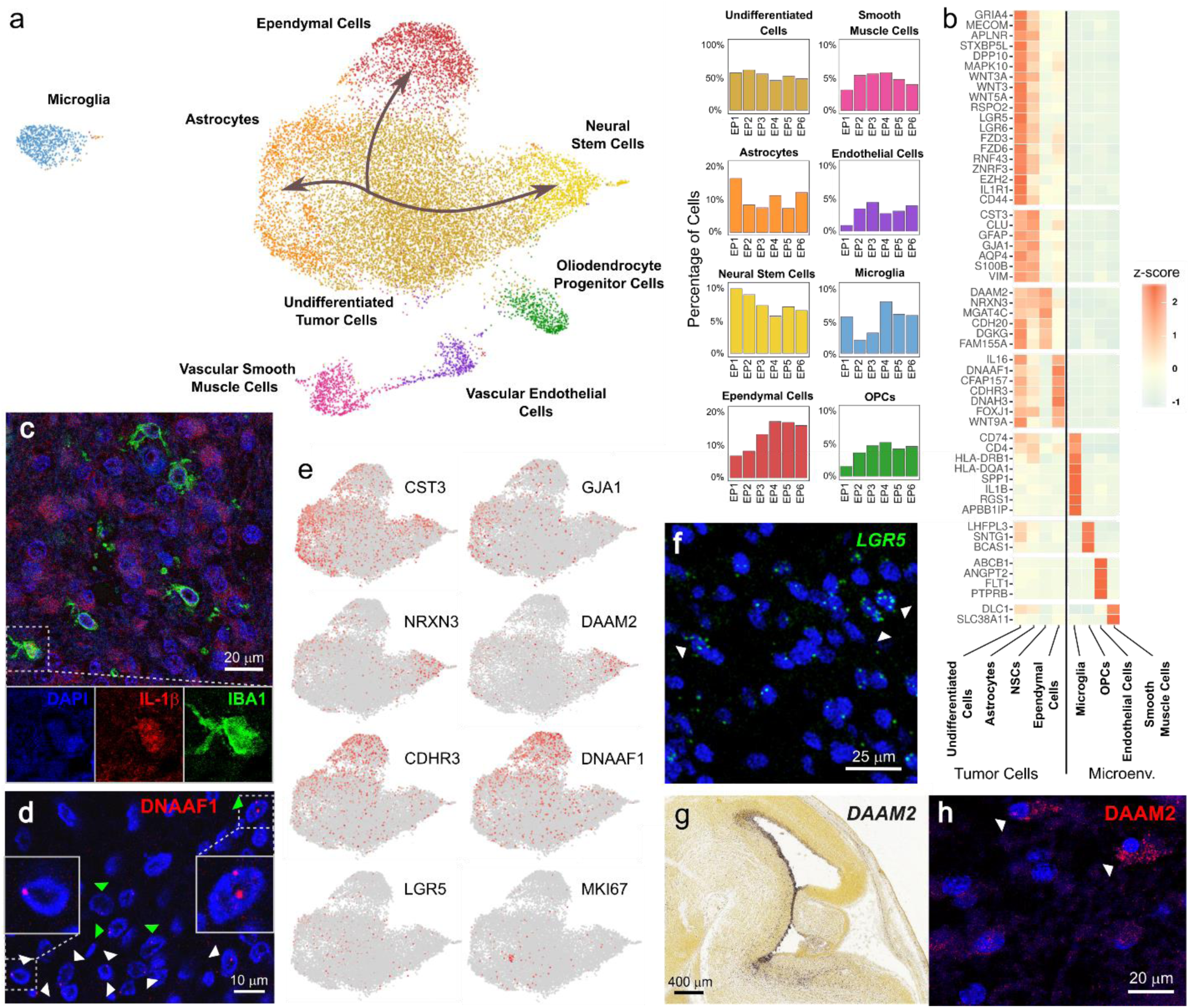
Single-nuclei RNA-seq of primary PFA tumors uncovers LGR5+ tumor stem cells, abundant WNT/β-catenin signaling, and a pro-inflammatory microenvironment. **a)** Single-nuclei RNA-seq data of 16,084 cells from 6 primary tumors. The UMAP representation of the data is labeled by the 8 cell populations identified. The three lineages of tumor-derived cells are indicated by arrows on the malignant cell population. The proportions of each cell type in each tumor is depicted in the bar plots. **b)** Some of the top differentially expressed genes in each population. Many of the genes expressed by tumor-derived cell populations are also expressed by undifferentiated tumor cells. Tumor cells express multiple genes coding for components of the WNT signaling pathway. **c)** Immunofluorescence staining of a primary PFA ependymoma (EP3) showing abundance of IL-1β (red) in the tumor microenvironment and the location of microglia marked by expression of IBA1. The detail in the bottom panel shows the localization of IL-1β in the cytoplasm of a tumor-infiltrating microglia. **d)** Immunofluorescence staining of a primary PFA ependymoma (EP3) for DNAAF1. The white (green) arrows indicate cells with one (two) foci of DNAAF1 protein localization. In the insets, two cells are shown in higher detail. **e)** The part of the UMAP corresponding to malignant cells is colored by the expression level of several genes that depict a significant gradient of expression (Laplacian score *q*-value < 0.01) along the cell differentiation lineages of the tumor. At the center is a population of cells with scattered expression of *LGR5* and upregulation of cell proliferation genes like *MKI67*, consistent with being tumor stem cells. **f)** Single-molecule RNA FISH showing the expression of *LGR5* in a primary PFA ependymoma (EP3). The arrows indicate a small number of cells with high expression levels of *LGR5* which, based on our single-nuclei transcriptomic analysis, would correspond to tumor stem cells. **g)** *In situ* hybridization for *DAAM2* expression in a sagittal section of the E15.5 mouse midbrain and hindbrain (Image credit: Allen Institute). **h)** Immunofluorescence staining of a primary PFA ependymoma (EP3) for DAAM2. The white arrows indicate two tumor-derived neural stem cells expressing DAAM2. Blue: DAPI.

### Primary PFA ependymal tumors promote a proinflammatory microenvironment

We first built a combined representation of the single-nuclei data of the 6 primary tumors (Fig. 1a and Supplementary Fig. 1). Upon clustering cells according to their expression profile and performing differential expression analysis, we identified 8 cell populations which we annotated according to the expression of known marker genes. These populations included undifferentiated tumor cells, ependymal cells, astrocytes, vascular endothelial cells, vascular smooth muscle cells, oligodendrocyte progenitor cells, microglia, and a population of *GFAP*^+^ *GJA1*^-^ *FOSJ1*^-^ *NRXN3*^+^ cells that we identified as neural stem cells based on the co-expression of astrocytic and neural genes (Fig. 1a,b and Supplementary Table 2). Each cell population was similarly represented across the 6 tumors, with undifferentiated tumor cells and microglia comprising approximately 50% and 5% of the cells respectively (Fig. 1a). Tumor-infiltrating microglia expressed MHC class II-related molecules characteristic of M1 glioma-associated microglia (Zeiner et al., 2015), including *CD4, CD74*, and MHC-II α/β chains, reflecting a proinflammatory microenvironment (Fig. 1b and Supplementary Fig. 2). Their expression profile also included osteopontin (*SPP1*), *APOE*, and *LPL* expression (Supplementary Fig. 2), which have been associated with active microglia in early postnatal neurogenic niches (Li et al., 2019) and neurodegenerative lesions (Masuda et al., 2019). To identify potential mechanisms of paracrine communication and chemotaxis between microglia and other tumor cell populations, we systematically searched for differentially expressed genes coding for ligand and receptor pairs (Supplementary Fig. 3 and Supplementary Table 3). We found that microglia expressed interleukin-1β (IL-1β) (Fig. 1b), a potent reactivator of the astrocytic hypoxia-angiogenesis program (Argaw et al., 2006), whereas the IL-1β receptor gene (*IL1R1*) was expressed by tumor cells (Fig. 1b). Conversely, tumor cells expressed high levels of IL-16 (Fig. 1b), a potent chemotactic cytokine that binds to the CD4 receptor (Center et al., 1996), and CD44, which interacts with microglial osteopontin to promote a glioma stem-cell-like phenotype (Pietras et al., 2014). We confirmed by immunofluorescence the abundance of the IL-1β protein in the tumor microenvironment and its cytoplasmic localization in tumor-infiltrating microglia (Fig. 1c). Taken together, these results suggest that primary PFA tumors promote a pro-inflammatory, microglia-rich, tumor microenvironment through abundant IL-16, IL-1β, and osteopontin signaling.

**Figure 2.**
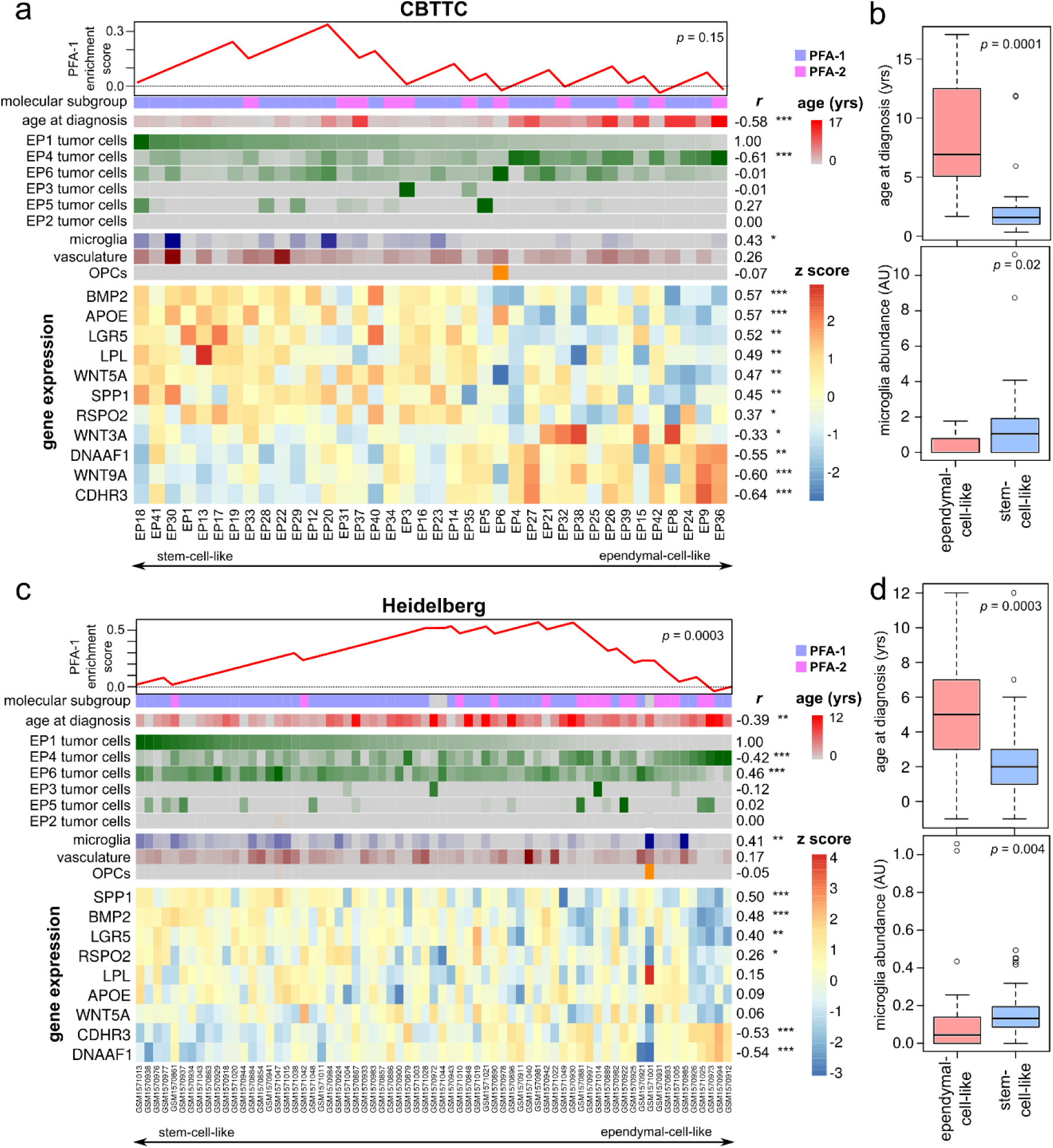
Stem-cell- and ependymal-cell-like PFA tumors have distinct expression profile, age of onset, and microenvironment composition. **a)** The single-nuclei gene expression signatures of PFA tumors EP1-6 were used to infer cell population abundances in the CBTTC cohort, comprising 38 pediatric PFA tumors profiled with bulk RNA-seq. Patients are arranged from left to right by decreasing inferred abundance of stem-cell-like (EP1-like) tumor cells. From top to bottom, the PFA-1 enrichment score, age of diagnosis, inferred abundance of tumor cells with distinct gene expression profile, inferred abundances of normal cell populations, and expression of representative genes are shown. On the right side, the Spearman’s correlation coefficient of each of these features with the inferred abundance of stem-cell-like tumor cells and its level of significance are indicated (test of association adjusted for multiple hypotheses testing using Benjamini-Hochberg procedure; *: *q*-value ≤ 0.05, ** *q*-value ≤ 0.01, ***: *q*-value ≤ 0.001). **b)** Tumors in the CBTTC cohort were classified as ependymal-cell-like (ependymal-cell-like tumor cell abundance > stem-cell-like tumor cell abundance) or stem-cell-like (ependymal-cell-like tumor cell abundance ≤ stem-cell-like tumor cell abundance). The age at diagnosis (top) and inferred abundance of microglia (bottom) are shown. **c)** and **d)** The same analysis as in a) and b) is presented for a cohort of 69 pediatric PFA tumors profiled with microarrays for gene expression by (Pajtler et al., 2015), showing consistency of the results.

**Figure 3.**
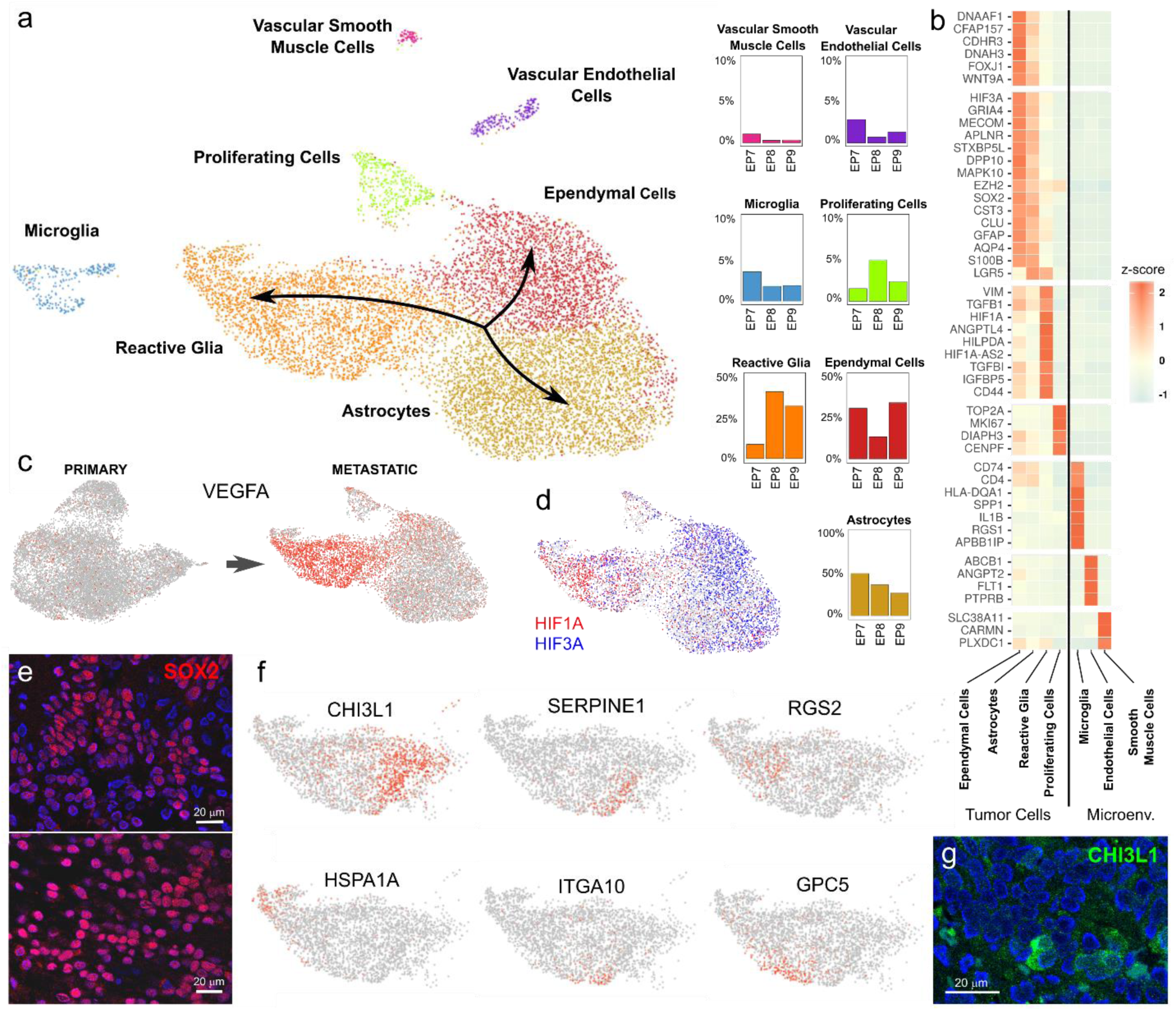
Metastatic progression of PFA ependymoma is associated with an EMT characterized by expression of hypoxic and reactive gliosis programs. **a)** Single-nuclei RNA-seq data of 10,640 cells from 3 metastases. The UMAP representation of the data is labeled by the 7 cell populations identified. The three lineages of tumor-derived cells are indicated by arrows. The proportions of each cell type in each tumor is depicted in the bar plots. **b)** Some of the top differentially expressed genes in each population. Tumor-derived reactive-glia have a mesenchymal phenotype and express high levels of hypoxia- and angiogenesis-related genes. **c)** The part of the UMAP corresponding to malignant cells in the primary and metastatic PFA tumors is colored by the expression level of *VEGFA*, showing the emergence of a population of *VEGFA*-expressing tumor reactive glia in metastatic tumors. **d)** The part of the UMAP corresponding to malignant cells in metastatic tumors is colored by the expression of *HIF1A* (red) and *HIF3A* (blue). The transition into reactive gliosis is associated with a switch between HIF3α and HIF1α expression. **e)** Immunofluorescence staining of a PFA ependymoma metastasis (top, EP9) and a primary PFA tumor (bottom, EP3) for SOX2 (red), showing an abundance of SOX2^low^ and SOX2^-^ cells in the metastasis as compared to the primary. Blue: DAPI. **f)** An analysis of the expression profile of tumor-derived reactive glia reveals much heterogeneity in this cell population. The part of the UMAP representation corresponding to this cell population is colored by the expression level of several genes that depict a significant localization of expression within that cluster (Laplacian score *q*-value < 0.01). **g)** Immunofluorescence staining of a PFA ependymoma metastasis (EP9) showing the presence of CHI3L1^+^ cells (green). Blue: DAPI.

### PFA ependymoma tumor cells recapitulate neurodevelopmental programs driven by WNT signaling

During normal development, ependymal cells, neural stem cells, and astrocytes differentiate from radial glia in neurogenic niches (Coletti et al., 2018; Ortiz-Alvarez et al., 2019; Redmond et al., 2019). In our analysis, these 3 cell populations appeared adjacent to the cluster of undifferentiated tumor cells in the expression space (Fig. 1a). They had high expression levels of genes upregulated by the undifferentiated tumor cell population, including PFA markers like *GRIA4, MECOM*, and *APLNR* (Pajtler et al., 2015; Taylor et al., 2005), as well as *DPP10, STXBP5L*, and *MAPK10* (Fig 1b; Benjamini-Hochberg *q*-value < 0.05), consistent with these populations representing tumor cells that have undergone partial cell differentiation (Jessa et al., 2019; Vladoiu et al., 2019). The proportions of tumor-derived neural stem cells and ependymal cells were strongly anticorrelated in our data (Pearson’s *r* = −0.96, *p*-value = 0.001, Supplementary Fig. 4), potentially reflecting differences in the homeostasis between neural stem cell and ependymal cell generation across patients. Consistent with this hypothesis, the ratio between the proportions of tumor-derived ependymal cells and neural stem cells was correlated with the ratio between the expression level of the DNA replication regulators Geminin and GemC1 (Spearman’s *r* = 0.77, *p*-value = 0.10), which respectively induce neurogenesis and ependymogenesis in radial glia of the adult neurogenic niche (Ortiz-Alvarez et al., 2019). Immunofluorescence for DNAAF1 (Fig. 1b), an axonemal dynein pre-assembly protein expressed by tumor-derived ependymal cells and localized at basal bodies (Basten et al., 2013), revealed a mixture of ependymal cells with predominantly one or two DNAAF1 foci per cell (Fig. 1d), consistent with the previously reported ciliary structure of ependymal tumors (Alfaro-Cervello et al., 2015).

**Figure 4.**
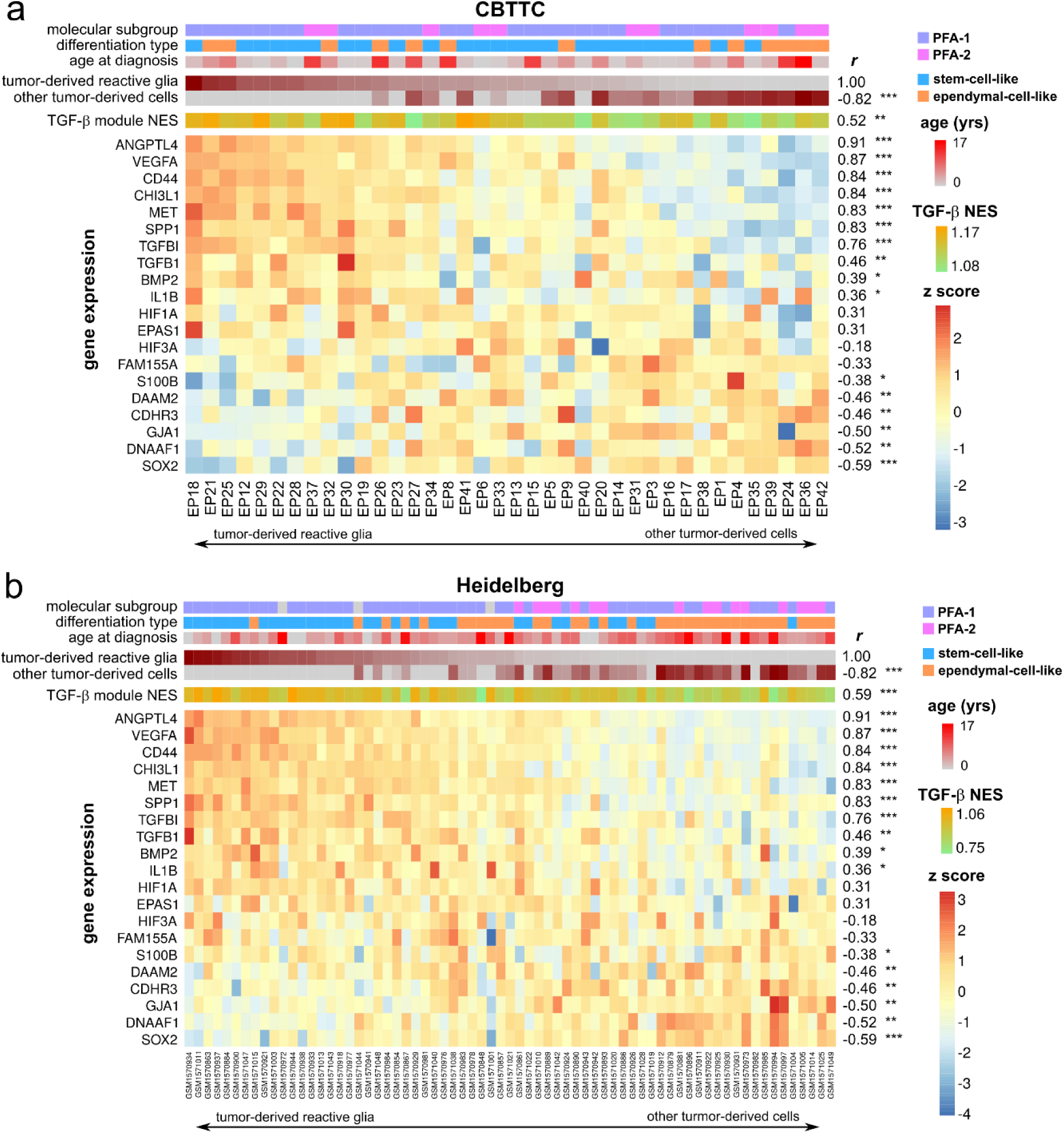
Mesenchymal PFA ependymoma cells are associated with abundant TGF-β signaling and stem-cell-like tumors. **a)** The single-nuclei gene expression signatures of PFA metastases EP7-9 were used to infer cell population abundances in 38 pediatric PFA tumors from the CBTTC cohort profiled with bulk RNA-seq. Patients are arranged from left to right by decreasing inferred abundance of reactive glia tumor cells. From top to bottom, the molecular subgroup, differentiation type, age of diagnosis, inferred abundance of reactive glia and other tumor cells, normalized enrichment score (NES) of a curated module of 189 genes upregulated by TGFB1, and expression of representative genes from the single-nuclei RNA-seq data analysis. On the right side, the Spearman’s correlation coefficient of each of these features with the inferred abundance of reactive glia tumor cells and its level of significance are indicated (test of association adjusted for multiple hypotheses testing using Benjamini-Hochberg procedure; *: *q*-value ≤ 0.05, ** *q*-value ≤ 0.01, ***: *q*-value ≤ 0.001). **b)** The same analysis as in a) is presented for a cohort of 69 pediatric PFA tumors profiled with microarrays for gene expression by (Pajtler et al., 2015), showing consistency of the results.

To identify gradients of expression related to cell differentiation, we used a spectral graph method (Govek et al., 2019). This approach identified 2,655 genes with significant patterns of expression along ependymal, astrocytic, or neuronal maturation trajectories (permutation test, Benjamini-Hochberg *q*-value < 0.05; Supplementary Table 4) and uncovered a proliferative subpopulation within the undifferentiated tumor cells that expressed high levels of *LGR5* (Fig. 1e), a canonical stem cell marker in multiple tissues and cancer types (Leung et al., 2018). The presence of this subpopulation was confirmed by single-molecule RNA fluorescent *in situ* hybridization (FISH) (Fig. 1f). Binding of R-spondin to LGR5, and/or its homologs LGR4 and LGR6, results in the stabilization of Frizzled receptors and the consequent amplification of canonical β-catenin-dependent WNT signaling. Consistently, we observed significant upregulation of genes coding for components of the WNT signaling pathway in the tumor cells (Fig. 1b and Supplementary Table 2), including expression of *RSPO2, WNT9A, WNT5A, WNT3A*, and *WNT3*, and Frizzled receptors, as previously observed at the bulk level (Palm et al., 2009). Additionally, tumor-derived neural stem cells expressed *DAAM2* (Fig. 1b,e), a modulator of canonical WNT signaling through PIP5K-PIP_2_ (Lee et al., 2015; Lee and Deneen, 2012), which is required for early dorsal neural stem cell specification in the developing spinal cord (Lee and Deneen, 2012) and is expressed in the ventricular zone of the prepontine hindbrain and the midbrain tegmentum of E15.5 mice (Fig. 1g). We confirmed by immunofluorescence the expression of the DAAM2 protein in a small subset of the tumor cells (Fig. 1h), in concordance with the results of our single-nuclei RNA-seq analysis. Taken together, these results highlight the importance of β-catenin-dependent WNT signaling programs in the proliferation and differentiation of PFA ependymoma tumor stem cells.

### Ependymal PFA tumors can be stratified according to their degree of differentiation

We next sought to identify gene expression differences among the 6 primary tumors that might be informative at the population level. To that end, we used our single-nuclei data to build a gene expression signature matrix consisting of differentially expressed genes between the transformed cells of each tumor, microglia, oligodendrocyte progenitor cells, and vascular cells (Supplementary Fig. 5). We used this matrix to infer the abundance (Newman et al., 2015) of these cell populations in the 38 PFA tumors for which bulk RNA-seq data was available. Most tumors in this cohort had an abundance of cells with an expression profile similar to that of transformed cells from two of the tumors (EP1 and EP4) (Fig. 2a). EP1-like tumors had a stem-cell-like gene expression signature, with high levels of *LGR5, RSPO2, WNT5A*, and *BMP2*, whereas EP4-like tumors had high expression of differentiated ependymal cell markers, including *DNAAF1* and *CDHR3*, as well as *WNT3A* and *WNT9A* (Fig. 2a). These two distinct signatures were strongly anti-correlated across the cohort (Spearman’s *r* = −0.61, *p*-value = 10^−4^) and separated PFA tumors into stem-cell- (*n* = 25) and ependymal-cell-like (*n* = 13) tumors (Fig. 2a). A gene set enrichment analysis (GSEA) for a general tissue stem-cell gene module (Wong et al., 2008) confirmed the association of stem-cell-like tumors with stem-cell gene expression programs (Spearman’s correlation between EP1 tumor cell abundance and GSEA normalized score *r* = 0.34, *p*-value = 0.04, Supplementary Fig. 6). Stem-cell-like tumors had a higher abundance of tumor-infiltrating microglia (fold-change 5.5, Wilcoxon rank-sum *p*-value = 0.02, Fig 2a,b), higher expression of pro-inflammatory microglial genes (Fig. 2a), and were associated with younger patients (average age 2.5 years, as compared to 8 years for ependymal-cell-like tumors, Wilcoxon rank-rum *p*-value < 10^−4^, Fig. 2a,b). We did not observe any significant difference in cell proliferation and overall patient survival between these two groups (Supplementary Fig. 7), although the small cohort size might be a limiting factor. These results were confirmed in an independent cohort of 69 pediatric PFA tumors profiled for gene expression with microarrays (Pajtler et al., 2015) (Fig. 2c,d and Supplementary Fig. 6). In both datasets we observed a significant correlation between the inferred abundances of microglia and vascular cells (Fig. 2a,c; Spearman’s *r* = 0.36, *p*-value = 0.03 for the CBTTC cohort, and *r* = 0.40, *p*-value < 0.001 for the Heidelberg cohort), suggestive of the role of microglia in tumor vascularization (Arnold and Betsholtz, 2013). Based on these results, we concluded that ependymal PFA tumors are heterogeneous regarding their degree of differentiation, and that differentiated tumors are associated with older ages of onset.

**Figure 5.**
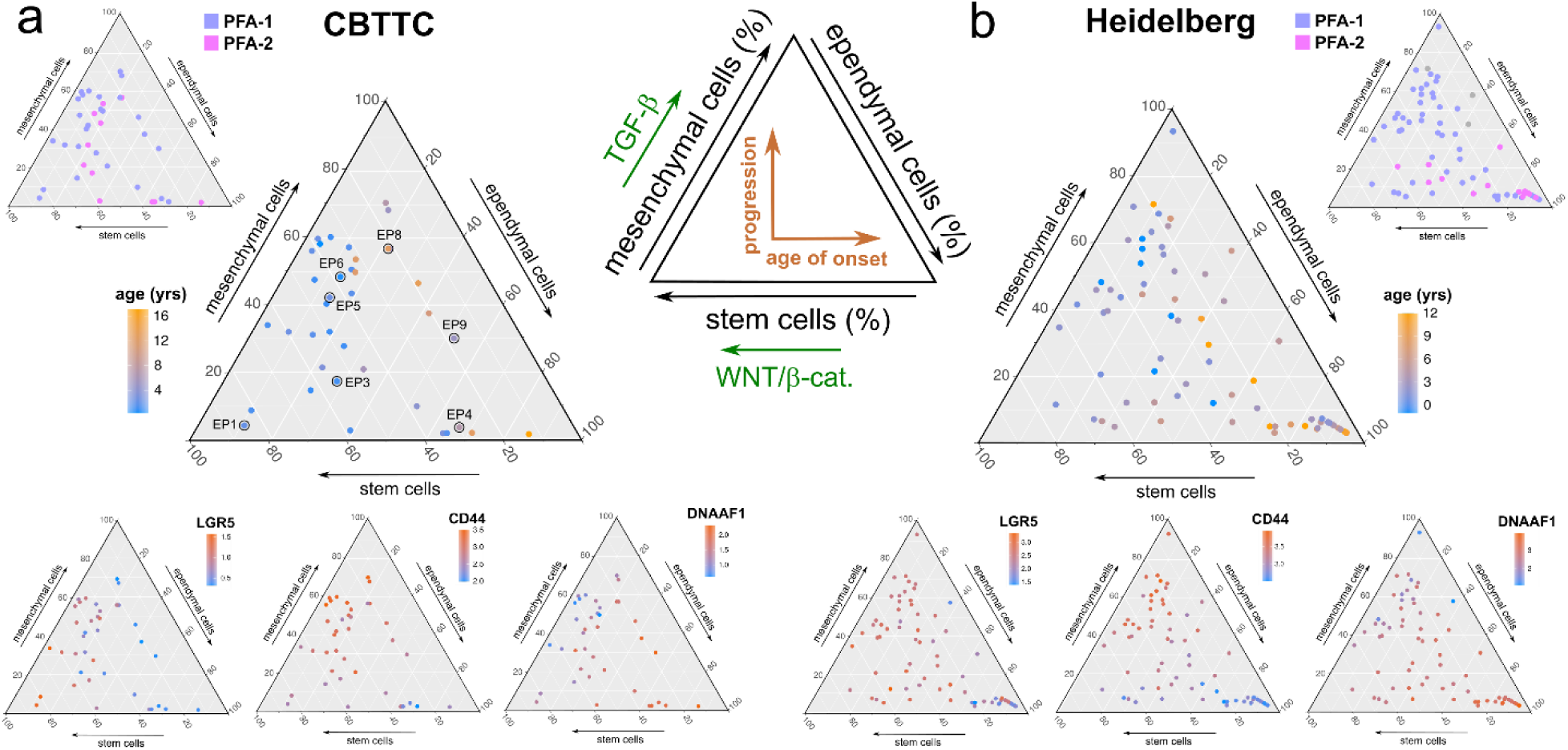
Stratification of pediatric PFA tumors according to their malignant cell type composition. **a)** Ternary plots depicting the 38 pediatric PFA tumors from the CBTTC cohort according to the proportions of tumor stem cells, ependymal cells, and reactive glia inferred from bulk RNA-seq data. The same ternary plot is colored by the tumor molecular subgroup (top left), age of diagnosis (center), and the gene expression level of representative markers for each of the three cell populations. The vertical and horizontal directions are respectively associated with tumor progression and age of onset. **b)** Same ternary plots for the 69 pediatric PFA tumors profiled with microarrays for gene expression by Pajtler et al., showing consistent results.

### PFA ependymoma metastases include a distinct mesenchymal cell population characterized by elevated expression of hypoxic and reactive gliosis programs

Having characterized the cell ecosystem and signaling pathways of primary tumors, we next studied the changes involved in metastatic tumor progression. To that end, we followed the same approach to analyze the 3 PFA metastases profiled with single-nuclei RNA-seq. Metastatic tumors recapitulated the differentiation trajectories of primary tumors with some remarkable differences (Fig. 3a,b and Supplementary Fig. 1). Consistent with our findings for primary tumors, we observed tumor-derived cells of varying degrees of differentiation, including ependymal cells, proliferative cells, reactive glia, and astrocytes (Fig. 3a,b). However, the gene expression profiles of tumor-derived cells were less distinct from each other than in primary tumors (Fig. 3b and Supplementary Table 5). Proliferating cells were abundant and clustered separately due to their strong cell cycle signature. Tumor cells expressed high levels of *LGR5* (Fig. 3a,b and Supplementary Table 5). Additionally, we did not observe a population of tumor-derived neural stem cells. Instead, metastatic tumors had a distinct *CD44*^high^ *TGFBI*^+^ cell population comprising ∼30% of the malignant cells (Fig. 3a,b and Supplementary Table 5). Expression of *CD44* and *TGFBI* has been associated with a TGFβ-driven mesenchymal phenotype in adult glioblastoma (Anido et al., 2010; Guo et al., 2018; Johansson et al., 2017; Pan et al., 2018) and with reactive astrocytes in brain lesions (Girgrah et al., 1991; Sherman and Back, 2008; Yun et al., 2002). Consistent with this mesenchymal reactive gliosis phenotype, *CD44*^high^ cells expressed high levels of hypoxia-angiogenesis genes, including *VEGFA, ANGPTL4, HIF1A, HIF1A-AS2, EPAS1*, and *HIGLPDA* (Argaw et al., 2012; Chakraborty et al., 2018; Mao et al., 2016; Mineo et al., 2016; Zhang et al., 2017), as well as the *MET* proto-oncogene (Jeon and Lee, 2017) (Fig. 3b,c and Supplementary Table 5; Benjamini-Hochberg *q*-value < 0.05). The transition from astroependymogenesis into reactive gliosis was associated with a switch between *HIF3A* and *HIF1A* expression and downregulation of the HIF3α target gene *SOX2* (Fig. 3b,d) (Forristal et al., 2010). Immunofluorescence staining for the SOX2 protein confirmed the abundance of SOX2^low^ and SOX2^-^ tumor cells in PFA metastases as compared to primary tumors (Fig. 3e). An analysis of tumor-derived reactive glia using the spectral graph method (Govek et al., 2019) revealed a great deal of heterogeneity within this cell population (Fig. 3f and Supplementary Table 6). This analysis uncovered multiple subpopulations of reactive glia characterized by the expression of *SERPINE1* and *CHI3L1*, which are also part of the mesenchymal gene expression signature in glioblastoma (Behnan et al., 2019), *ITGA10, GPC5, RGS2, HSPA1A*, and other genes (Fig. 3f and Supplementary Table 6). Immunofluorescence staining for CHI3L1 confirmed the presence of CHI3L1 expressing cells in PFA metastases (Fig. 3g). Taken together, our results suggest that metastatic progression of pediatric PFA ependymoma is associated with the emergence of a distinct mesenchymal population of *CD44*^high^ *SOX2*^low^ cells in the tumor that expresses hypoxic, angiogenic, and reactive gliosis gene expression programs with high plasticity.

### The mesenchymal PFA ependymoma gene expression program is associated with abundant TGF-β signaling and stem-cell-like tumors

To verify the results of our single-nuclei transcriptomic analysis of metastatic tumors in the larger cohort of PFA tumors, we used the single-nuclei RNA-seq data to build a gene expression signature matrix that included tumor-derived reactive glia (Supplementary Fig. 8). We used this matrix to infer the abundance of this cell population in each of the 38 PFA tumors profiled with bulk RNA-seq. Consistent with the single-nuclei analysis, the abundance of tumor-derived reactive glia was strongly correlated with the expression of hypoxia- and angiogenesis-related genes (Fig. 4a). Additionally, tumors with a high abundance of tumor-derived reactive glia expressed high levels of pro-inflammatory cytokines, such as osteopontin and IL-1β (Fig. 4a). *TGFB1* and TGF-β induced genes (Padua et al., 2008) were upregulated in these tumors (Fig. 4a, Spearman correlation between the TGF-β module GSEA score and reactive glia abundance *r* = 0.53, *p*-value < 0.001, median GSEA score *q*-value = 0.003), adding support to a possible link between the mesenchymal phenotype of PFA ependymoma and TGF-β signaling. Consistent with this hypothesis, the mesenchymal gene expression signature of glioblastoma multiforme (Neftel et al., 2019; Verhaak et al., 2010) was also enriched in PFA ependymal tumors with a high abundance of tumor-derived reactive glia (Supplementary Fig. 9, Spearman correlation between mesenchymal glioblastoma module GSEA score and reactive glia abundance *r* = 0.88, *p*-value = 2 × 10^−13^, median GSEA score *q*-value = 5 × 10^−10^). Our analysis also identified a significant inverse relation between the abundance of tumor-derived reactive glia and the expression of *SOX2* and ependymal, astrocytic, and neuronal differentiation markers at the bulk level (Fig. 4a), in agreement with the transition between reactive gliosis and astroependymogenesis observed in the single-nuclei RNA-seq analysis. Thus, stem-cell-like PFA ependymal tumors were also associated with a higher abundance of tumor-derived reactive glia (Fig. 4a, Spearman’s correlation between the abundances of tumor-derived reactive glia and EP1 tumor cells *r* = 0.34, *p*-value = 0.04). All of the statistical associations were confirmed in an independent cohort of 69 pediatric PFA tumors (Pajtler et al., 2015) (Fig. 4b and Supplementary Fig. 9). Our analysis of bulk transcriptomes is therefore consistent with the existence of a pediatric PFA ependymoma mesenchymal gene expression signature associated with TGF-β signaling and a cell population of tumor-derived mesenchymal reactive glia.

## Discussion

Tumor cell composition is the result of multiple concurrent molecular processes. Here, we have performed single-nuclei RNA-seq of primary and metastatic pediatric PFA ependymal tumors to investigate their cell composition and signaling pathways across tumor progression. Our study has identified two major processes that contribute to shaping the cell ecosystem of this cancer. On one side, tumor stem cells differentiate into multiple tumor-derived cell lineages that recapitulate the developmental trajectories of radial glia in neurogenic niches. Our data indicates that WNT/β-catenin signaling is a key regulator of this process and suggests a dorsal neurodevelopmental origin for these tumors. On the other side, tumor stem cells acquire a mesenchymal phenotype upon tumor progression. This mesenchymal phenotype is characterized by abundant TGF-β signaling and the expression of hypoxia- and angiogenesis-related programs, and it is reminiscent of the reactive astrogliosis programs that take place during brain injury and neurodegeneration. Our data indicates that the transition of PFA ependymoma tumor stem cells into this mesenchymal state is a discrete transition, characterized by upregulation of CD44 and HIF1A, and downregulation of SOX2. This transition might be favored by the proinflammatory cytokine milieu promoted by tumor cells in cooperation with infiltrating microglia. Thus, the progression of PFA ependymal tumors into a mesenchymal phenotype have many resemblances with the epithelial-mesenchymal transition of glioblastoma (Iwadate, 2016).

Our single-nuclei transcriptomic analysis therefore suggests that it is possible to stratify pediatric PFA ependymal tumors according to their degree of cell differentiation and mesenchymal progression (Fig. 5). We have identified gene expression signatures and markers for these two cell processes and have used these signatures to stratify 107 pediatric PFA ependymal tumors from two different cohorts based on their bulk transcriptomic profile. Our analysis uncovered an association of stem-cell-like PFA tumors with younger patients. It is currently unclear whether this association is due to differences in the neurodevelopmental time and cell of origin of these tumors, or to cell growth differences that lead to an earlier detection.

During the final stages of this work, two papers with related and partially overlapping results have appeared (Gojo et al., 2020; Michealraj et al., 2020). Based on a series of experiments involving *in vitro* culture of patient-derived tumor cells under hypoxic conditions, Michealraj et al. (Michealraj et al., 2020) found that hypoxic programs are important drivers of pediatric PFA ependymoma tumorigenesis, and related these programs to the hypoxic environment of the early embryo. On the other hand, Gojo et al. (Gojo et al., 2020) performed single-cell RNA-seq of a large cohort of pediatric ependymal tumors across multiple molecular groups and found the same tumor-derived cell differentiation trajectories for PFA ependymal tumors. However, in this work they did not consider metastases nor the interaction with the tumor microenvironment. Our work therefore provides an independent confirmation of the results of these two studies and complements them by investigating the relation between metastatic PFA ependymoma tumor progression, the expression of hypoxia-angiogenesis programs, the epithelial-mesenchymal transition of PFA tumor stem cells, and the proinflammatory microenvironment of PFA ependymoma. Our results highlight the importance of WNT/β-catenin, TGF-β signaling, and microglia-secreted cues for regulating pediatric PFA ependymoma tumor stem cell proliferation and metastatic progression, therefore contributing to the identification of candidate therapeutic targets for treating this cancer.

## Acknowledgements

The authors are grateful to the CBTTC for providing data and tissue specimens for conducting this study. They also thank Peng Hu, Qi Qiu, and Hao Wu for assistance with the implementation of sNuc-Drop-seq, Yugong Ho and Stephen Liebhaber for assistance with general experimental aspects, Doug Epstein for useful scientific discussions, and the Next-Generation Sequencing Core of the University of Pennsylvania for assistance with cDNA library sequencing. This work has been partially supported by an Advisory Council Research Award from the CBTTC. The work of R. A. is supported by the NHGRI T32 Computational Genomics Training Grant, NIH T32HG00046.

## Author contributions

R. A. carried out the computational analyses. E. C. T. performed the experiments. A. N. A. assisted with some of the computational analyses. M. S. and M. P. N. provided histopathology expertise. R. A., E. C. T., and P. G. C. wrote the manuscript. P.G.C. conceived and supervised the study.

## Methods

### Tumor samples

De-identified flash-frozen tumor specimens and formalin-fixed paraffin-embedded (FFPE) slides were provided by the CBTTC biorepository (CBTTC Approved Biospecimen Project 29). The anatomic location of the tumors and their diagnosis were obtained from the de-identified surgical, radiology, and pathology reports. All procedures were performed according to the institutional regulations of the University of Pennsylvania and the Children’s Hospital of Philadelphia (CHOP). The PFA identity of the tumors was confirmed based on the expression of PFA markers in bulk RNA-seq data (see *Bulk RNA-seq data processing*). For specimens with no available bulk RNA-seq data (EP2 and EP7), the expression of PFA markers was assessed by quantitative reverse transcription PCR. First strand cDNA synthesis was performed using Maxima H Minus Reverse Transcriptase (Thermo Scientific, cat# EP0753) according to the manufacturer’s protocol, with Oligo(dT)18 (Fisher Scientific, cat# SO131). cDNA concentration was measured with a Qubit 3 Fluorometer (Life Technologies). We used 10 ng of cDNA for each RT-qPCR reaction and KiCqStart SYBR Green primer pairs for GAPDH, L1CAM, APLNR, IFT46, CXorf67, and MECOM (Sigma Aldrich). SYBR FAST Universal qPCR Master Mix with low Rox (Roche Sequencing, cat# KK4602) was used and the reaction was performed with a QuantStudio 7 Flex (Applied Biosystems). Four technical replicates of each sample-primer combination were performed and used to determine the standard error.

### Bulk RNA-seq data processing

De-identified raw RNA-seq data of 40 pediatric ependymal tumors located in the posterior fossa were provided by the CBTTC biorepository (Felmeister et al., 2016) (CBTTC Approved Data Project 19). Gene expression was quantified for each tumor using Kallisto (version 0.45.1) (Bray et al., 2016) with the human reference genome GRCh38. To assign each of the 40 tumors a molecular group, we developed a classification method based on gene set enrichment analysis (GSEA) (https://github.com/CamaraLab/EPN_Classifier). This method determines whether a gene signature *s* is significantly enriched among a ranked list of genes *L* ordered by expression. Gene signatures *s* were constructed from an independent cohort of ependymal tumors profiled for expression by microarrays (Pajtler et al., 2015). For each of the eight molecular groups in this cohort (supratentorial sub-ependymoma, supratentorial ependymoma with YAP1-fusion, supratentorial ependymoma with RELA-fusion, posterior fossa sub-ependymoma, posterior fossa ependymoma group A, posterior fossa ependymoma group B, spinal myxopapillary ependymoma, and spinal anaplastic ependymoma), we used limma’s linear model (version 3.34.1) (Phipson et al., 2016) on this data to identify the top *n* = 50 differentially expressed genes exclusively upregulated in each group (*p*-value < 0.01 and log2 fold change > 2). Additionally, we utilized the gene signatures for PFA-1 and PFA-2 tumors from the bulk transcriptomic data of (Pajtler et al., 2018), consisting of the top *n* = 58 upregulated genes exclusive to each signature. To assess the importance of each signature on a query bulk RNA-seq dataset, we calculated a running-sum statistic for each signature *s* by walking down the complete list *L* of genes in the query dataset in decreasing order of expression. For gene *j* in *L*, the statistic increases by 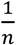 if *L*_*j*_ ∈ *s* or decreases by 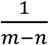 if *L*_*j*_ ∉ *s*, with *m* the total number of genes in *L*. By randomizing the genes in *s*, we used a permutation test to estimate the statistical significance of each score. If the p-value of all signatures was above 0.35, no molecular group was assigned. The training error of our classifier was 1.4% (3/209). To further benchmark the performance of our classifier, we ran it on an extended cohort of 94 pediatric ependymal tumors of mixed anatomical origin from the CBTTC biorepository for which bulk RNA-seq data were available. The classifier correctly predicted the anatomic location of the tumors in 92.6% (87/94) of the cases.

### Single-nuclei RNA-seq library preparation and sequencing

Single-nuclei RNA-seq was performed as previously described (Hu et al., 2017) with several modifications. Approximately 2 mm^3^ of tissue was cut from each tumor. Samples were incubated at room temperature in 1 ml homogenization buffer for 3 minutes, then dissociated with 13 strokes of a tight pestle in a 2 ml glass homogenizer. Samples were then filtered with a 40 μm mesh filter followed by a 30 μm mesh filter. After brief tabletop centrifugation, the nuclei pellet was resuspended in 1 ml PBA-BSA 0.01% and counted. Nuclei were defined by shape, size, and general appearance, and diluted to 100 nuclei/μl. Single-nuclei RNA-seq library preparation was performed as previously described (Hu et al., 2017) using a Drop-seq microfluidic system. cDNA quality of the libraries was evaluated with a bioanalyzer and quantified using a KAPA Library Preparation Kit (Roche sequencing, KK4824). Libraries were sequenced using an Illumina NextSeq 500 on high output mode with 20 bp (read 1) and 60 bp (read 2) paired end reads.

### Single-nuclei RNA-seq data processing

The Drop-seq computational pipeline (Macosko et al., 2015) was used to map to the human reference genome (GRCh38) via STAR alignment (version 2.6.1a) (Dobin et al., 2013), correct the cellular and molecular barcodes for sequencing errors, and build a count matrix. To account for poly-adenylated pre-mRNAs in the nucleus, we included reads that aligned to intronic regions of the genome (locus_function_list = intronic). We removed low-quality cells, debris, and empty droplets based on the inflexion point in a plot of the number of UMIs in each droplet ranked by decreasing order of magnitude. Additionally, we plotted the number of UMIs (in log scale) against the proportion of mitochondrial genes for each droplet. For most samples, the distribution of droplets in this scatter plot was bimodal. We used a linear cut to segregate cells (high number of UMIs and low percentage of mitochondrial genes) from debris (high percentage of mitochondrial genes and low number of UMIs) in this plot. We then followed the Seurat pipeline (version 3.0.2) (Satija et al., 2015) for normalizing, clustering, and visualizing the single-nuclei RNA-seq data. Upon log-normalizing and centering the expression data, we used Seurat’s anchor method (Stuart et al., 2019) to build combined representations for the primary (Fig. 1a) and metastatic tumors (Fig. 3a). We clustered the data using Louvain community detection based on the top 20 principal components and used Uniform Manifold Approximation and Projection (UMAP) (Becht et al., 2018) to construct 2-dimensional visualizations.

### Differential expression analysis

We used edgeR’s general linear model (version 3.20.1) (Robinson et al., 2010) to identify differentially expressed genes between each cell cluster and all the other cells in the single-nuclei RNA-seq data. We used the R package iTALK (Wang et al., 2019) to annotate differentially expressed genes (edgeR *q*-value < 0.01) coding for ligand and receptor pairs. For each cluster, we only considered genes expressed in at least 10 cells and with a log2 fold change greater than 1.5. Apart from these analyses, we used the R package RayleighSelection (Govek et al., 2019) to compute the Laplacian score on the log-normalized expression matrix and identify genes with a significant gradient of expression within the undifferentiated tumor cell (for primary tumors) and the reactive glia clusters (for metastatic tumors). We used Pearson’s correlation distance as metric and took the radius parameter ε of RayleighSelection to be the median pairwise distance among cells. We considered genes expressed in 1%-10% of the cells for the undifferentiated cell population, and 3-40% of the cells for the reactive glia population. To reduce the runtime, we used a random sample of 2,000 cells in each cluster.

### Gene set enrichment analysis

We used the R packages msigdbr (Liberzon et al., 2011; Subramanian et al., 2005) and fgsea (Korotkevich et al., 2019) to download the gene modules TGFB_UP.V1_UP (Padua et al., 2008) and WONG_ADULT_TISSUE_STEM_MODULE (Wong et al., 2008) from the MSigDB database and compute GSEA normalized enrichment scores and *p*-values in each bulk RNA-seq dataset. In addition, we considered the mesenchymal glioblastoma gene module from (Neftel et al., 2019).

### Deconvolution of bulk RNA-seq data

We built gene expression signature matrices by aggregating the TPM normalized single-nuclei RNA-seq data over each cell type of interest. Only genes that were differentially expressed (edgeR p-value < 0.01) in exactly one cell type were included in the signature matrix. To improve the specificity of expression signatures, we filtered out genes expressed in less than 5% of cells for a given cell type or those with < 2 fold change in expression. We used CIBERSORTx (Newman et al., 2019) to infer the absolute abundance of these cell populations in the bulk transcriptomic data (absolute mode and zero permutations).

### Immunofluorescence

Formalin-fixed paraffin-embedded tumor slides were deparaffinized in preparation for immunofluorescence with xylene. Washes of decreasing ethanol percentage were used to rehydrate tissue, and heat-induced epitope retrieval was performed using a universal antigen retrieval reagent (Novus Biologicals CTS015-NOV) for 10 minutes. After blocking with 1% horse serum in PBS, tissue was incubated with primary antibodies overnight at 4°C at the following dilutions: goat anti-IBA1 1:500 (ab178846, Abcam), rabbit anti-DAAM2 1:50 (HPA054630, Sigma Aldrich), rabbit anti-IL1B 1:100 (ab2105, Abcam), rabbit anti-DNAAF1 1:200 (HPA074239, Sigma Aldrich), mouse anti-SOX2 1:50 (sc-365823, Santa Cruz Biotechnology), and rabbit anti-CHI3L1 1:100 (12036-1-AP, Proteintech). Secondary antibodies were incubated for 1 hour at room temperature at the following dilutions: anti-goat Alexa 488 1:500 (a11055, Invitrogen), anti-rabbit Alexa 647 1:500 (a150075, Invitrogen), and anti-mouse cy5 1:500. Images were acquired using a Leica TCS SP8 Multiphoton Confocal Microscope at 40x magnification.

### Single-molecule RNA fluorescence *in situ* hybridization

LGR5 single molecule RNA FISH was carried out using the RNAscope Multiplex Fluorescent Reagent Kit v2 Assay (323100, Advanced Cell Diagnostics) using standard conditions and manual target retrieval. The *LGR5* probe (311021, Advanced Cell Diagnostics) was detected using Opal 520 (FP1487001KT, Akoya Biosciences) at a 1:1500 dilution. Images were acquired using a Leica TCS SP8 Multiphoton Confocal Microscope at 40x magnification.

