## Supplementary Figures and Supplementary Table Captions for "Cell Ecosystem and Signaling Pathways of Primary and Metastatic Pediatric Posterior Fossa Ependymoma"

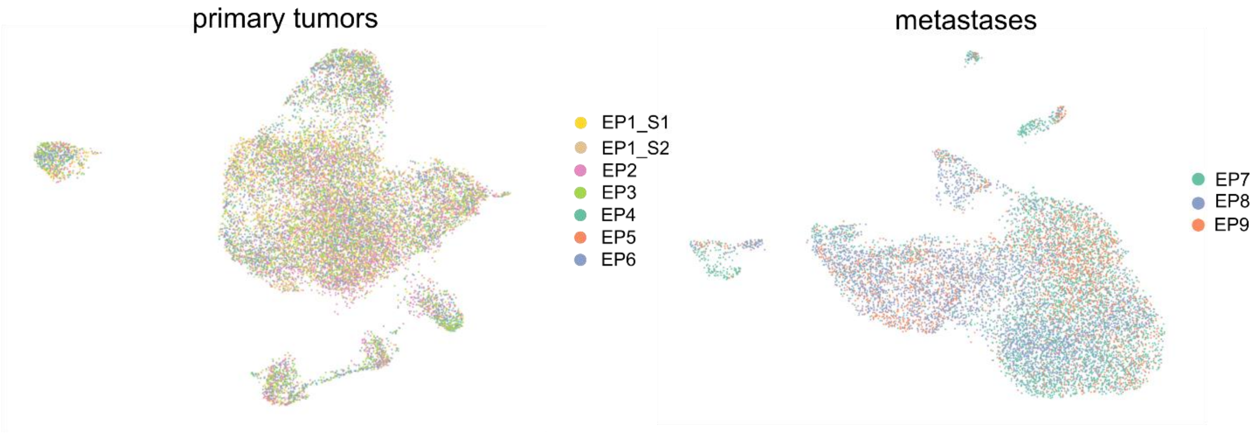


**Supplementary Figure 1. UMAP representations of the single-nuclei RNA-seq data of primary and metastatic PFA ependymal tumors colored by the sample of origin.** EP1_S1 and EP1_S2 represent two different samples from the same tumor specimen.

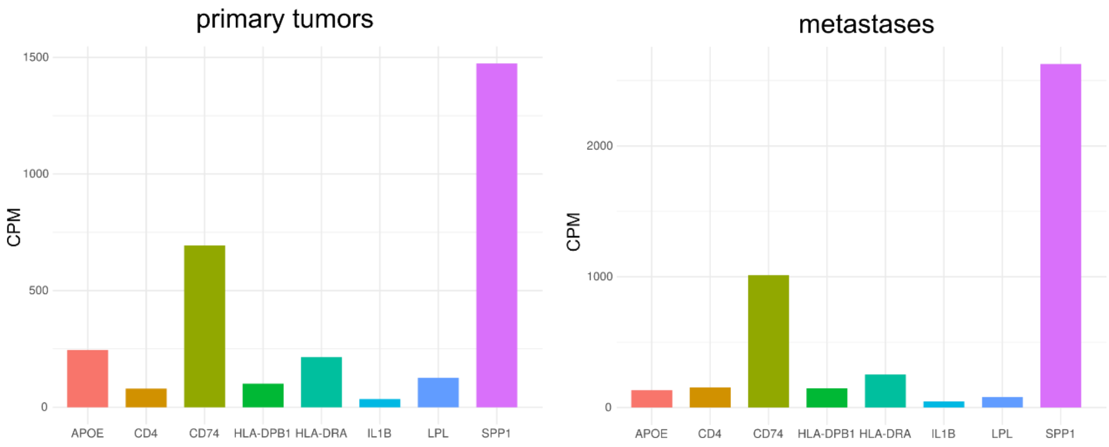


**Supplementary Figure 2. Expression of selected genes in microglia from the PFA ependymoma primary tumors and metastases.** The expression level of each gene is shown in counts per million (CPM) based on the single-nuclei RNA-seq data.

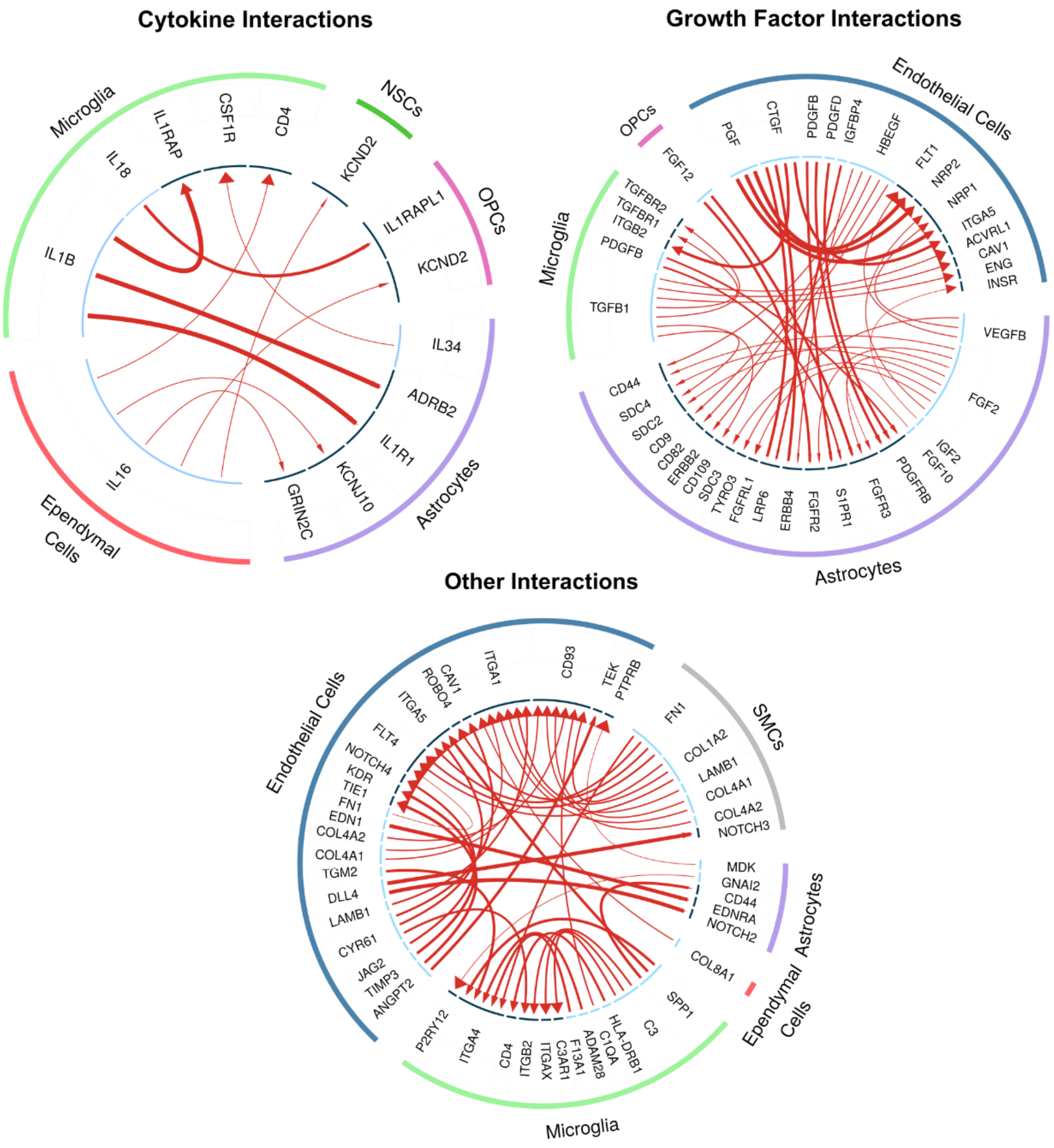


**Supplementary Figure 3. Differentially expressed genes (*q* < 0.01) coding for pairs of ligands and receptors in primary PFA tumors.** For other interaction only the top 40 significant pairs are shown. The complete list of differentially expressed pairs can be found in Supplementary Table 3. SMCs: smooth muscle cells. OPCs: oligodendrocyte progenitor cells. NSCs: neural stem cells.

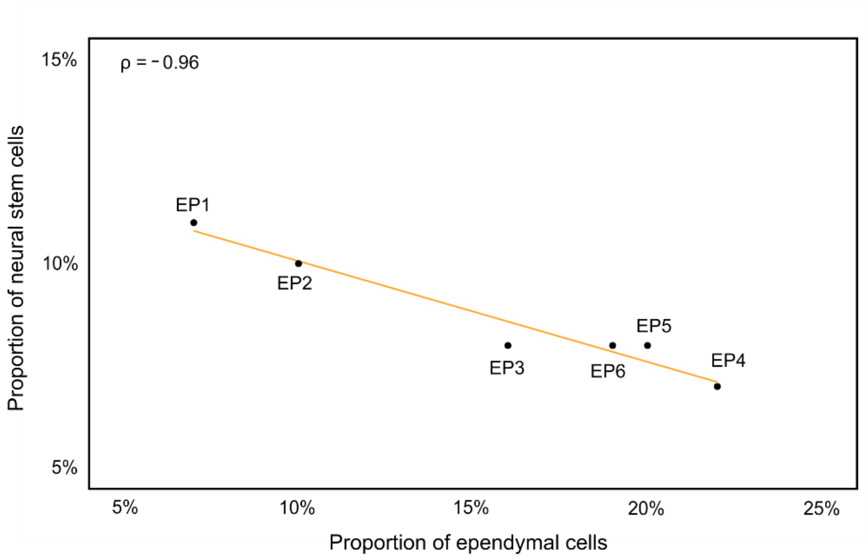


**Supplementary Figure 4. Relation between the proportion of tumor-derived neural stem cells and ependymal cells in each of the tumors profiled by single-nuclei RNA-seq.**

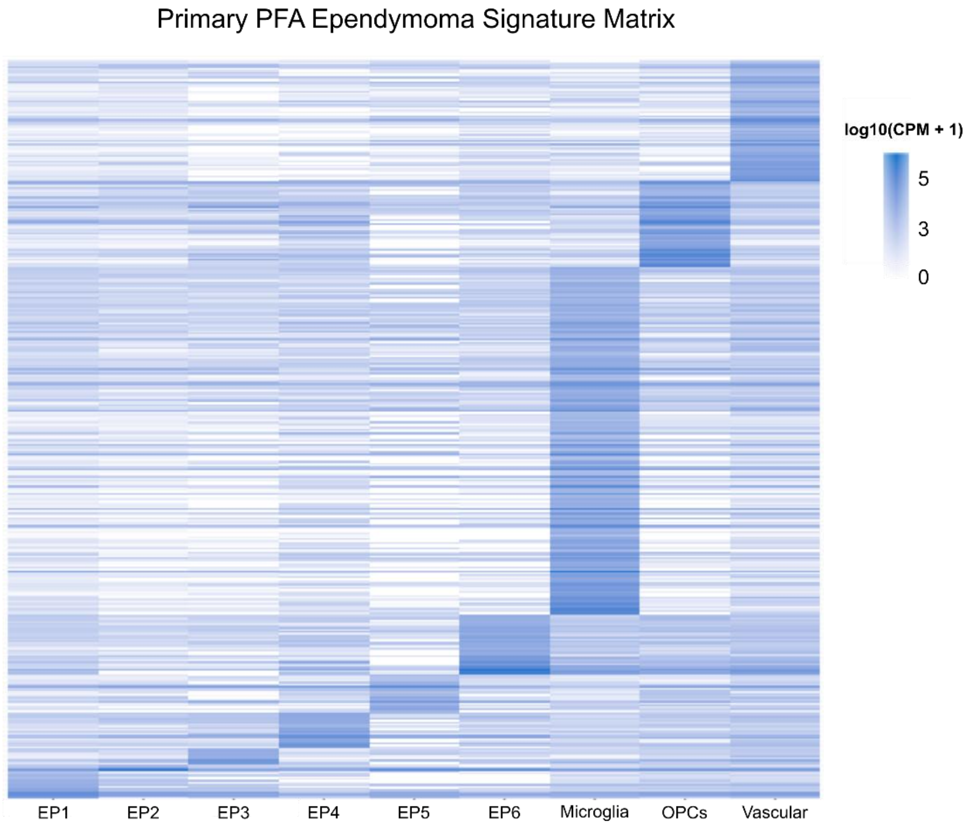


**Supplementary Figure 5. Gene expression signature matrix from the primary PFA ependymal tumors profiled by singe-nuclei RNA-seq.**

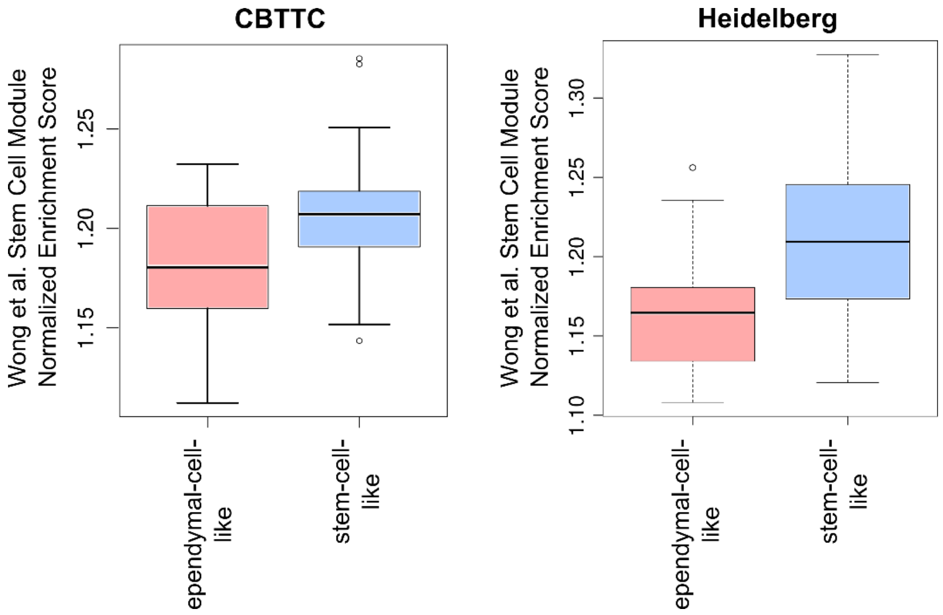


**Supplementary Figure 6. Gene set enrichment scores for the stem-cell gene expression module of (Wong et al., 2008) in stem-cell- and ependymal-cell-like PFA ependymal tumors.** The enrichment scores are shown for tumors profiled with bulk RNA-seq in the CBTTC cohort (left, Spearman's correlation between EP1 tumor cell abundance and normalized enrichment score $r = 0.34$, $p$-value = 0.04; median GSEA score Benjamini-Hochberg $q$-value = $10^{-9}$) and with expression microarrays in the Heidelberg cohort (right, Spearman's correlation between EP1 tumor cell abundance and normalized enrichment score $r = 0.52$, $p$-value < $10^{-6}$; median GSEA score Benjamini-Hochberg $q$-value = $2\times10^{-5}$). Box-plot elements: center line, median; box limits, upper and lower quartiles; whiskers, 1.5× interquartile range; points, outliers.

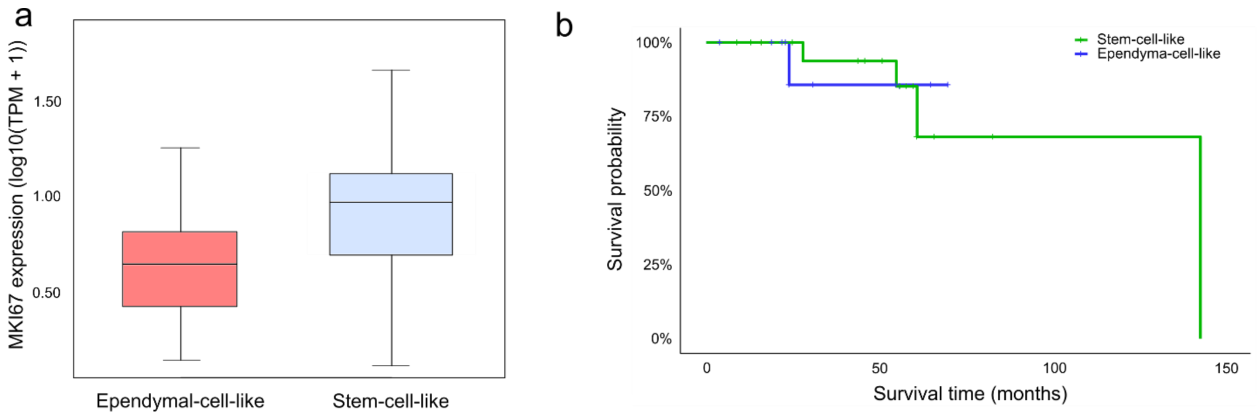


**Supplementary Figure 7. Comparison of cell proliferation and patient survival for 38 ependymal-cell- and stem-cell-like PFA tumors from the CBTTC cohort profiled by bulk RNA-seq. a)** Gene expression levels of the MKI67 cell proliferation marker (mean fold change = 1.3, two-sided Wilcoxon rank sum *p*-value = 0.09). **b)** Kaplan-Meier survival curves for ependumal-cell- and stem-cell-like tumors (log-rank test, *p*-value = 0.99).

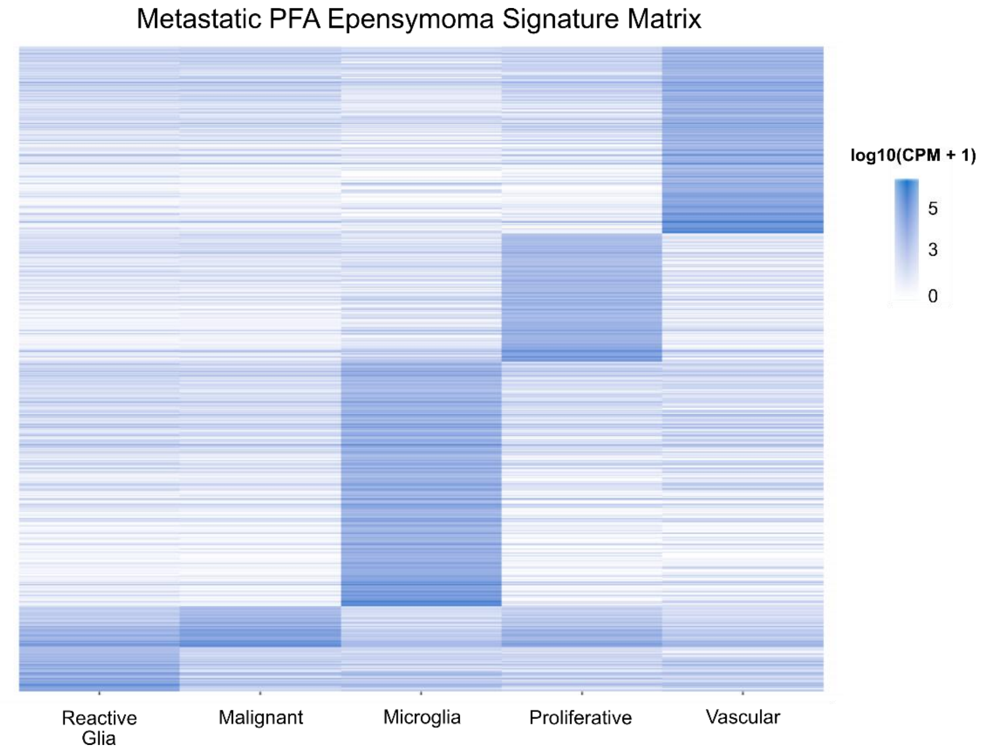


**Supplementary Figure 8. Gene expression signature matrix from the metastatic PFA ependymal tumors profiled by singe-nuclei RNA-seq.**

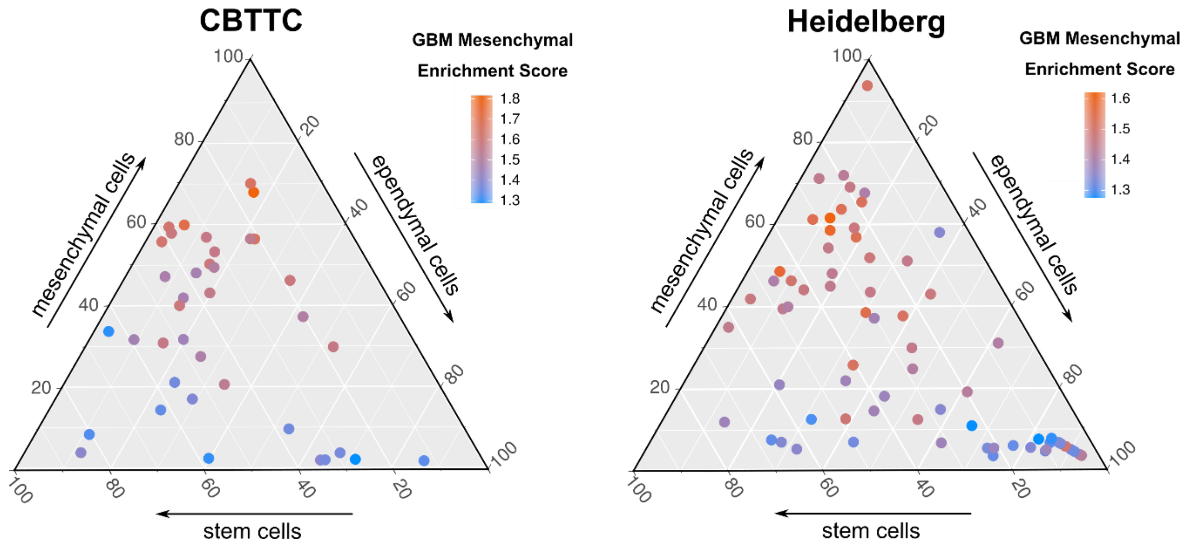


**Supplementary Figure 9. Mesenchymal pediatric PFA tumors are enriched for the mesenchymal gene expression signature of glioblastoma.** Ternary plots depicting the pediatric PFA tumors from the CBTTC (left) and Heidelberg cohorts (right) according to the proportions of tumor stem cells, ependymal cells, and reactive glia inferred from bulk RNA-seq data. The plots are colored by the normalized enrichment score of the mesenchymal gene expression signature of glioblastoma (median GSEA score Benjamini-Hochberg *q*-value = $5 \times 10^{-10}$ for the CBTTC cohort and $6 \times 10^{-6}$ for the Heidelberg cohort).

### Supplementary Tables

**Supplementary Table 1. Summary of the patients considered in the study.** Provided as a separate file.

**Supplementary Table 2. Differentially expressed genes in each cell population for the primary PFA ependymal tumors profiled by single-nuclei RNA-seq.** Provided as a separate file.

**Supplementary Table 3. Differentially expressed genes coding for pairs of ligands and receptors for the primary PFA ependymal tumors profiled by single-nuclei RNA-seq.** Provided as a separate file.

**Supplementary Table 4. List of genes with significant Laplacian score in the expression space of the undifferentiated tumor cell population for the primary PFA ependymal tumors profiled by single-nuclei RNA-seq.** Provided as a separate file.

**Supplementary Table 5. Differentially expressed genes in each cell population for the metastatic PFA ependymal tumors profiled by single-nuclei RNA-seq.** Provided as a separate file.

**Supplementary Table 6. List of genes with significant Laplacian score in the expression space of the reactive glia cell population for the metastatic PFA ependymal tumors profiled by single-nuclei RNA-seq.** Provided as a separate file.
